# Differential reach vector computations in mIPS and PMd as revealed through HD-tDCS

**DOI:** 10.1101/2022.11.22.517546

**Authors:** Sisi Xu, Jason Gallivan, Gunnar Blohm

## Abstract

Recent neural and behavioural findings provide support that the medial intraparietal sulcus (mIPS) and dorsal premotor (PMd) activity reflect aspects of a kinematic plan for reaching movements. However, it is unclear how these two regions differentially contribute to reach planning. Here, we used high-definition transcranial direct current stimulation (HD-tDCS; 4 x 1 electrode placement; 2 mA for 20 min; 3 cm radius) to investigate the functional roles of mIPS and PMd in the left hemisphere of humans. We examined the changes in endpoint error in reaching task with different initial hand positions and different target locations spanning both visual hemi-fields. Participants completed the task with (stimulation, post-stimulation) and without stimulation (pre-stimulation) of individually fMRI-localized cortical areas mIPS and PMd. We found a significant interaction effect between initial hand position (IHP) and target position on the difference in horizontal endpoint error after cathodal left mIPS stimulation and significant IHP and target position main effects after cathodal left PMd stimulation, suggesting that IHP and target position are not yet integrated into a movement vector at the input of the mIPS, but are integrated in the input of PMd. Hence, these findings reveal a distinction between mIPS and PMd in the stages of movement vector formation for reaching movements and indicate that HD-tDCS is a viable method for perturbing localized cortical activity to elucidate localized cortical function.

## Introduction

To plan goal-directed reaches, we need to compute a movement plan that specifies the reach goal location relative to the initial hand position (Vindras & Viviani, 1998). The calculation of this difference vector is believed to occur during the transformation from the initial conditions of the hand and target position into the motor command (Batista, Buneo, Snyder, & Andersen, 1999), providing aspects of the movement plan, i.e. amplitude and direction information for movement execution (Krakauer, Pine, Ghilardi, & Ghez, 2000; Bock & Arnold, 1992). However, where and how spatial information is stored and combined in the brain persist as a gap in our understanding of this process.

A number of parietofrontal circuits are thought to be devoted to the process of planning for different movement types ahead of movement execution. One particular example is the dorsomedial pathway, which comprises regions relevant to reaching component of reach-to-grasp actions: the medial part of IPS (mIPS), the parieto-occipital junction (POJ), and the dorsal premotor cortex (PMd) (Jeannerod, Arbib, Rizzolatti, & Sakata, 1995). Studies of the early component of the dorsomedial pathway, the mIPS, using transcranial magnetic stimulation (TMS) tend to suggest a role in encoding visual information about the initial hand position from visual cortex (Vesia & Crawford, 2012) but has also been implicated in processing movement direction (Davare, Zénon, Desmurget, & Olivier, 2015). Thus, there is conflicting indications of a disposition upstream as opposed to downstream of the visuomotor transformation.

The later component of the dorsomedial pathway, the PMd, also plays a role in reach planning, based on delay period activity in monkey PMd which was thought to represent the movement plan for reaching movements as the target location selectivity was independent of choice of hand use (Cisek, 2002). Appearing to complementary to the direction coding role of the mIPS, the PMd has been suggested to be involved in amplitude coding (Davare, Zénon, Desmurget, & Olivier, 2015). Yet, potential differences between the two regions in the formation of the movement vector remain to be examined. What part of the movement plan (i.e., IHP, target position, or difference vector) is encoded in mIPS and PMd?

Transcranial direct current stimulation (tDCS) can serve as a valuable tool for non-invasive and reversible modulation of cortical excitability to unravel cortical structure-function relationships. A weak electrical direct current is applied to the scalp to induce polarity-dependent subthreshold modulations and consequent changes in cortical excitability in the underlying cortex. Anodal stimulation produces neuronal depolarization and promotes neuronal excitation in the stimulated cortical area while cathodal stimulation in hyperpolarization and neuronal inhibition (Bindman et al. 1962; Purpura & McMurtry, 1965). The high definition 4 x 1 configuration offers the ideal stimulation focality to study the function of a more specific cortical area. The 4×1 HD-tDCS configuration which consists of small disc electrodes arranged in a ring significantly enhances spatial accuracy and precision due to the smaller electrode size, ring radius (distance between centre and surround electrode), and the configuration-determined current paths (Datta, Truong, Minhas, Parra, & Bikson, 2012; Datta et al, 2009). The central electrode, which determines the polarity, is placed over the target and the 4 surrounding return electrodes are placed around it to guide current flow. It is not known how HD-tDCS of mIPS or PMd may affect movement planning. To our knowledge, HD-tDCS has not been applied previously in the study of motor planning.

Here, we stimulated the mIPS and PMd in the left hemisphere and examined participants’ performance on a right-hand reaching task and inferred the functional roles of the two areas based on performance changes. We hypothesized polarity-specific behavioral effects on the basis of polarity-specific neurophysiological effects, i.e., somatic depolarization (hyperpolarization) by anodal (cathodal) stimulation. Our results provide evidence that HD-tDCS can modulate performance in a movement planning task to reveal differences in the functional properties of mIPS and PMd.

## MATERIALS AND METHODS

### Participants

Ten volunteers (age range: 20 – 39; 7 females) participated in the experiment. Participants had normal or corrected-to-normal vision and were right handed. They performed two tasks with their dominant right hand: an fMRI localizer session followed by the main tDCS experiment. Participants provided written informed consent to the experimental conditions that were approved by the Queen’s University General Board of Ethics, in compliance with the Declaration of Helsinki.

### Overview of Experiment

Each participant underwent a localizer session consisting of one anatomical scan and one functional scan. Subsequently, all subjects made 4 visits for the HD-tDCS experiment (2 polarity x 2 sites), with a one-week interval between visits. (Note: experiments and analyses were carried out in 2014-2016)

### MRI Acquisition & Localizer Task

Imaging was conducted using a 3-Tesla Magnetom Trio scanner (Siemens Medical Systems, Erlangen, Germany). Participants rested supine in the scanner and wore a 32-channel head coil. At the beginning of the localizer session, a 176 slice, high-resolution T1-weighted anatomic reference volume was acquired using a 3D MP-RAGE sequence (single shot, ascending sequence in the sagittal plane with TR = 1760 ms, TE = 2.2 ms, FoV = 256 mm, flip angle = 9°, and voxel size = 1 mm^3^).

Participants were then scanned with blood oxygenation level-dependent (BOLD) fMRI for the localization of the reaching network during an interleaved center-out pointing and saccade task (T2*-weighted gradient echo planar imaging, FOV = 24.0 cm, matrix = 80 x 80, in-plane resolution of 3mm, TR = 2 s, TE = 30 ms, FA = 90°, 35 contiguous axial slices acquired in an interleaved order along the AC-PC line). Visual stimuli were rear-projected using an LCD projector (NEC LT 265 DLP projector; resolution, 1024 x 768; 60 Hz refresh rate) on a screen mounted behind the participant, and viewed by the participant via a mirror mounted to the head coil directly above the eyes. The visual presentation consisted of a circular arrangement of 8 white circles (~15 degrees of visual angle) surrounding a central red or green square (Figure 1). Participants were instructed to make self-paced centre-out pointing movements to each white circle in the circular arrangement while fixating on a central fixation square when it is red and execute centre-out saccades when the central square is green. The intertrial interval was 2 s. In each movement block (pointing and saccade conditions), participants started each respective centre-out movement towards the 12 o’clock white circle and proceeded in a clockwise manner until the end of the block. The reach-saccade localizer included 8 reaching blocks (16 s per block), 8 saccade blocks (16 s per block), and 3 fixation/baseline blocks (10 s per block), which were placed at the beginning, middle and end of the run. The localizer lasted 5 min 22 s (157 brain volumes).

**Figure 1:**
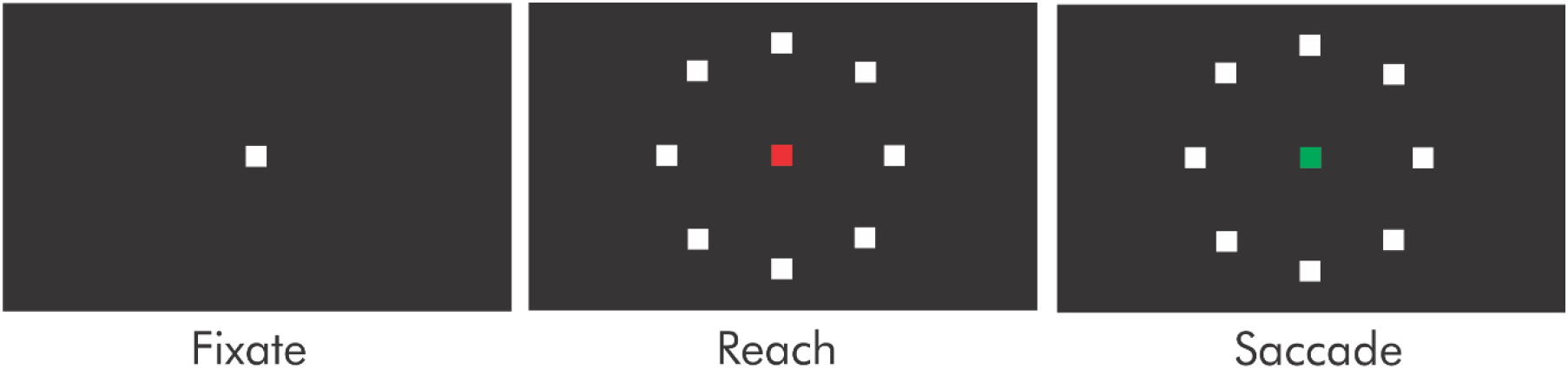
Functional localizer task display. Participants performed the localizer task while laying supine inside the MRI. The visual stimuli were rear-projected using an LCD projector. Participants were instructed to fixate when the white square appears, and either perform centre-out reaches with the right hand or saccades when the white square turns respectively, red or green. Participants started each respective centre-out movement towards 12 o’clock and proceeded in a clockwise manner until the end of the block.

The functional localizer data were superimposed on the corresponding participant’s high resolution anatomical image with an alignment on the AC-PC plane. All preprocessing and univariate analyses were performed using Brain Voyager QX version 2.6 (Brain Innovation). Preprocessing for the localizer data included slice scan-time correction, 3D motion correction (alignment of the first volume of the functional scan, which was closest in time, to the anatomical scan), high-pass temporal filtering of 3 cycles/run, functional-to-anatomical co-registration, and spatial smoothing (trilinear-sinc interpolation performed during realignment and sinc interpolation performed during reorientation). We contrasted BOLD signal for the pointing and fixation conditions rather than the pointing and saccade conditions to identify the hotspots of the left mIPS and left PMd due to known activation overlap in response to saccade and reach planning (Vesia et al., 2010). The left mIPS and left PMd were reliably identified in each participant by significantly higher response during the pointing condition compared to the fixation condition. The location of the left mIPS was defined by selecting the peak activity voxel or voxel cluster medial to the intraparietal sulcus in the left hemisphere (Vingerhoets, 2014). The location of the left PMd was defined by selecting the peak activity voxel or voxel cluster at the junction of the precentral sulcus and the superior frontal sulcus in the left hemisphere (Amiez, Kostopoulos, Champod, & Petrides, 2006).

### Co-registration

The scalp location corresponding to the target cortical area was identified using Brainsight (Rogue Research Inc., Montreal, Quebec) and marked to guide electrode placement so that the peak electric field of HD-tDCS would be directed at the targeted cortical area. The nasion, nose tip, left and right inter-tragal notches were used as landmarks to guide participant-image registration, and the targets (peak voxel representing the left mIPS and PMd) were selected as the foci for stimulation from the superimposed functional image data. The marked location was the minimum inline projection of the target mapped onto the scalp. The MNI coordinates of each intended site of stimulation were recorded for comparisons with those reported in the literature. The mean normalized MNI coordinates (±standard deviation [SD]) of the stimulation points were −25 ± 5, −58 ± 6, and 52 ± 6 mm for the left mIPS and −25 ± 6, −9 ± 5, and 59 ± 5 mm for the left PMd (x, y, z), consistent with the locations reported in studies using various approaches summarized in Table 1 (mIPS coordinates reported in fMRI studies: Davare, Zenon, Pourtois, Desmurget, & Olivier, 2012; Grefkes, Ritzl, Zilles, & Fink, 2004; Prado et al., 2005; Stark & Zohary, 2008; Blangero, Menz, McNamara, & Binkofski, 2009; mIPS coordinates used in TMS studies: Vesia, Prime, Yan, Sergio, & Crawford, 2010; Davare, Zenon, Pourtois, Desmurget, & Olivier, 2012; PMd: Meta-analysis of 31 studies, Mayka, Corcos, Leurgans, & Vaillancourt, 2006; PMd coordinates reported in fMRI study: Gallivan, McLean, Smith, & Culham, 2011).

**Table 1.**
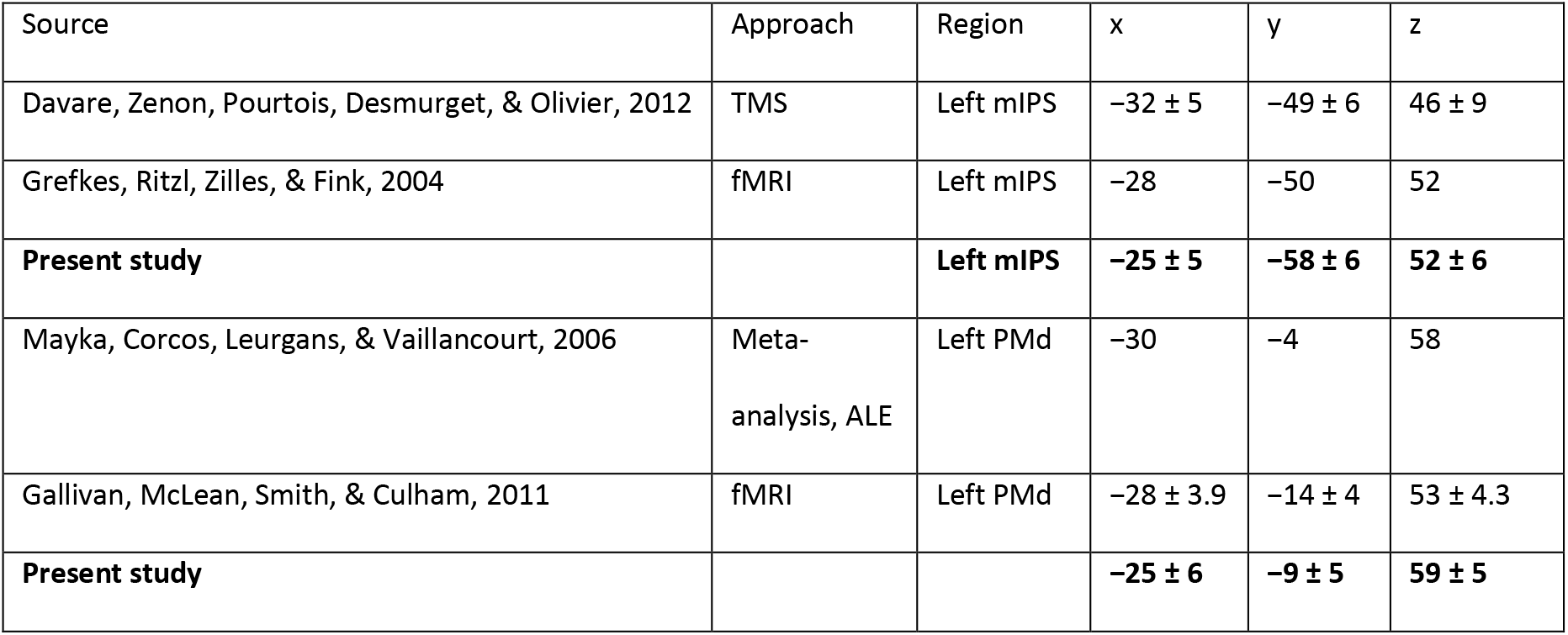
MNI coordinates reported in studies using different approaches.

### High-Definition Direct Current Stimulation

Continuous direct current was generated by a microprocessor-controlled constant current source (DC-STIMULATOR MC, NeuroConn GmbH, Ilmenau, Germany), and delivered using sintered Ag/AgCl ring electrodes (1cm diameter) for 20 min at a current strength of 2mA. The 4×1 electrode configuration was used and consisted of a central electrode placed on marked position from co-registration, and surrounded by four pick-up electrodes at a radius of 3 cm. To improve the conductance, hair was washed and dried prior to the experiment, and separated under the scalp using cotton swabs. Electrodes were stabilized within plastic holders filled with conducting gel embedded in a mesh cap which was fastened on the participant’s head with straps. The impedance was under 20 kΩ throughout stimulation.

On separate days, anodal or cathodal stimulation were administered to the left mIPS or the left PMd for a total of 4 sessions. At each site, the order of the stimulation polarity was randomly counterbalanced across participants. Each session consisted of three conditions, pre-stimulation (control), stimulation, and post-stimulation and each participant completed 250 trials per condition (50 trials per block x 5 blocks per condition), for a total of 750 trials per session (Figure 2).

**Figure 2.**
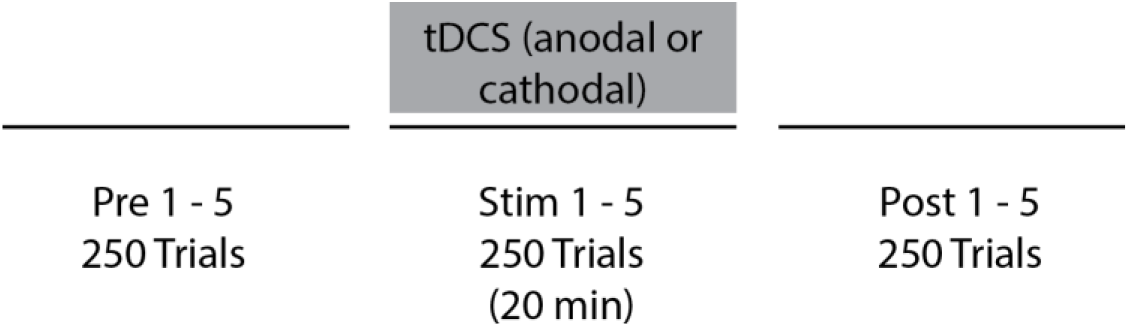
Schematic representation of the experimental protocol. Pre blocks 1 - 5 were collected prior to stimulation to measure the participant’s baseline performance for that day. For stim blocks 1 - 5, anodal or cathodal tDCS was applied to either the left mIPS or the left PMd for 20 min. After stimulation, post blocks 1-5 were collected to examine how tDCS-induced effects evolve over time. EEG activity was collected in pre blocks 1-5 and post blocks 1-5.

### Experimental Setup

Participants were seated in a dark room with their head immobilized in an upright position by dental impressions attached to a tilted reaching setup, such that their straight-ahead gaze aligned with a fixation cross (Figure 3). Visual stimuli (i.e., initial hand position, fixation cross, target) were presented on a tilted overhead monitor and viewed in a tilted semi-silvered mirror located midway between the tilted reaching surface and the monitor. All surfaces were tilted at 30°. Hand position was occluded by the semi-silvered mirror but was displayed as a representative visual marker to guide the initial alignment of the right index fingertip to the starting position and extinguished at movement onset.

**Figure 3.**
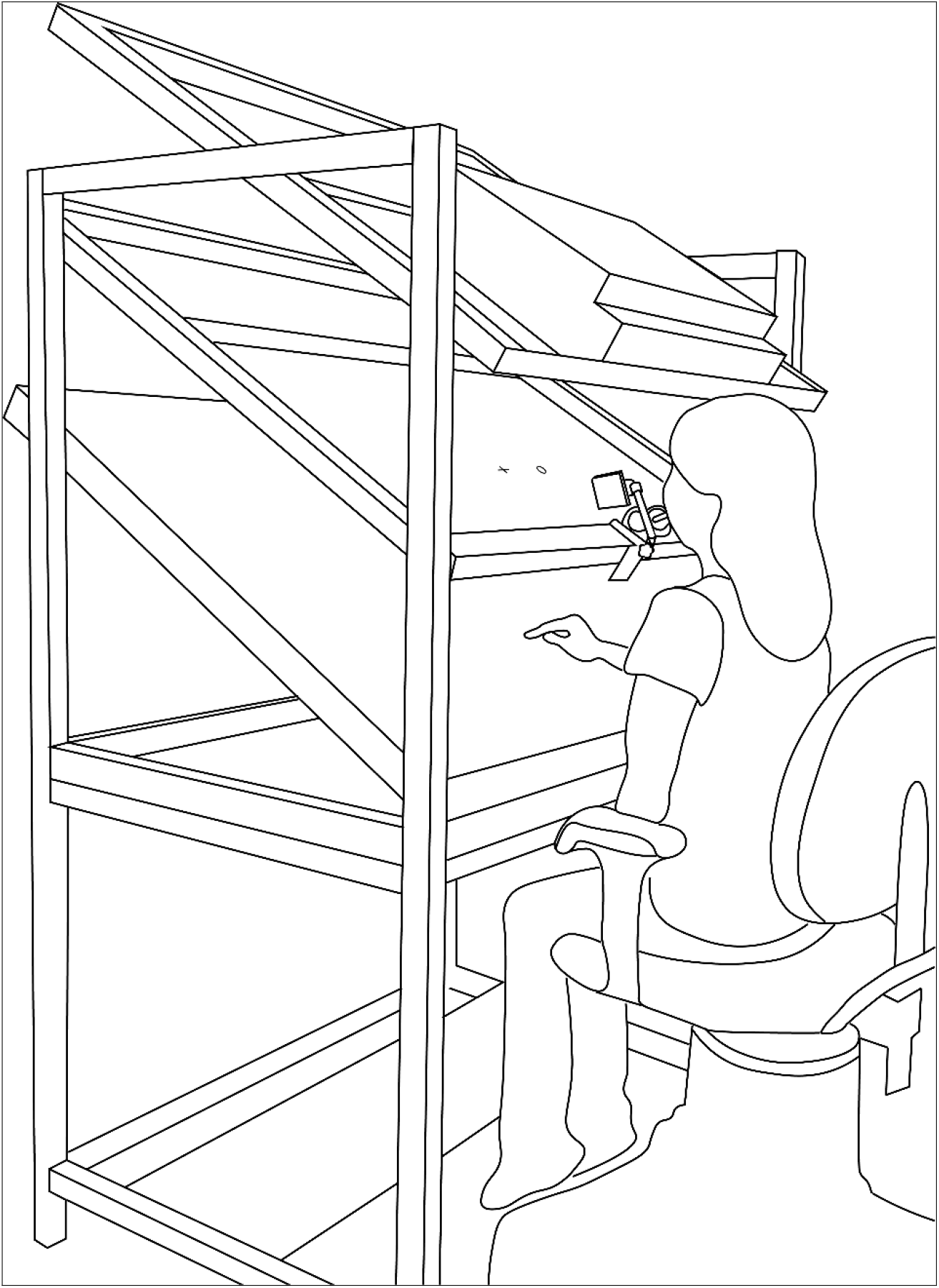
Schematic representation of experimental setup. Participant is seated upright in front of a virtual reality set-up with their head immobilized by an individual dental impression. Visual stimuli is presented on an overhead monitor and viewed in a semi-transparent mirror, which allowed the reflected visual stimuli to be in the plane of the reaching surface.

Eye fixation was monitored using EyeLink 1000 eye tracking system (SR Research; sampling rate, 500 Hz; accuracy, 0.5°). Hand movement data were obtained by localizing the 3D position of infrared light-emitting diodes taped to the tip of the right index finger (Optotrak Certus; Northern Digital; sampling rate, 400 Hz; accuracy, 0.1 mm.

We recorded a measure of cortical excitability in the targeted cortical regions using electroencephalography before and after stimulation. This set-up included V-Amp 16, OpenViBE driver, Vamp-16 EEG amplifier (Brain Products), sintered Ag/AgCl electrodes, a reference electrode attached to the skin surface above participant’s vertebrate C7 and a ground electrode attached to the elbow. This data was not used for the following analyses.

### Reaching Task

The reaching task required participants to make reaching movements from one of 2 initial hand positions (7.5cm left/right of midline) to one of 4 randomly chosen briefly flashed targets (20cm distant, 5cm apart horizontally) while fixating on a straight-ahead cross in-line with the targets. The trial began with the presentation of a marker to direct the participant to the initial hand position (IHP). Upon aligning the right index fingertip with the IHP, the IHP cue extinguished and a fixation cross appeared (500 – 1000 ms after trial start), during which a target was briefly presented. The participant was allowed to move immediately in response to the brief presentation of the target and was instructed to maintain their gaze on the fixation cross when the target flashed as well as during the reaching movement.

Reaching task display and timings are presented in Figure 4, and outlined above eye position (Figure 5B), hand position (Figure 5C) and velocity plots (Figure 5D) for an exemplary participant reach (Figure 5A). All target-directed reaches were performed without visual feedback of the hand. Instructions and practice (10 trials) were provided prior to the main task.

**Figure 4:**
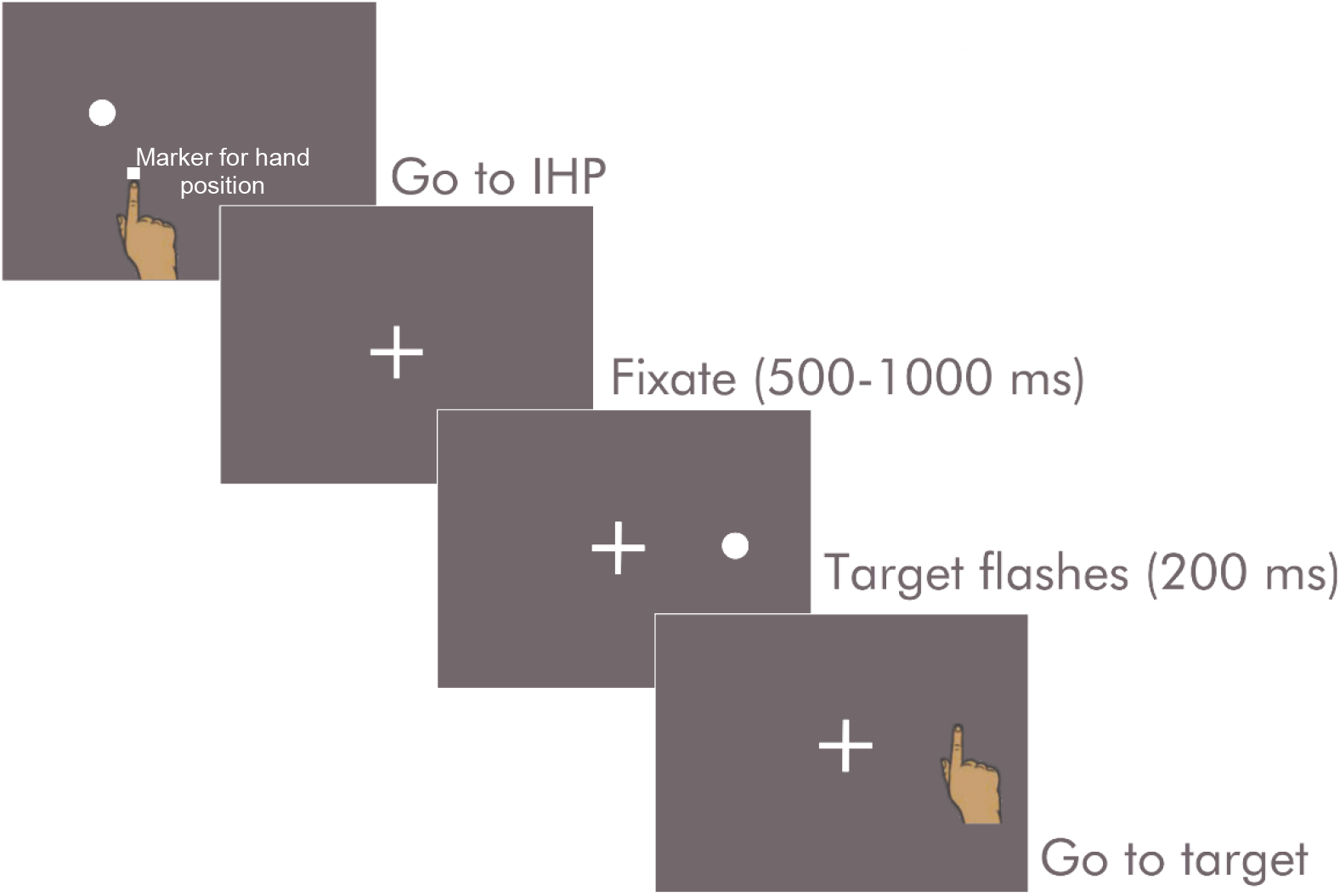
Reaching task display and timing. A typical trial begins with the participant aligning their right index finger (indicated by a visible marker) to the IHP cue. Vision of the hand is occluded by the semi-silvered mirror. However, the participant is provided with a marker representing the hand position prior to target-directed reaching. A fixation cross then appears, followed by the brief presentation of the target. The participant then moves immediately to the target and the marker indicating hand position is extinguished at the onset of movement. Target viewing and target-directed reaches are performed with peripheral vision.

**Figure 5:**
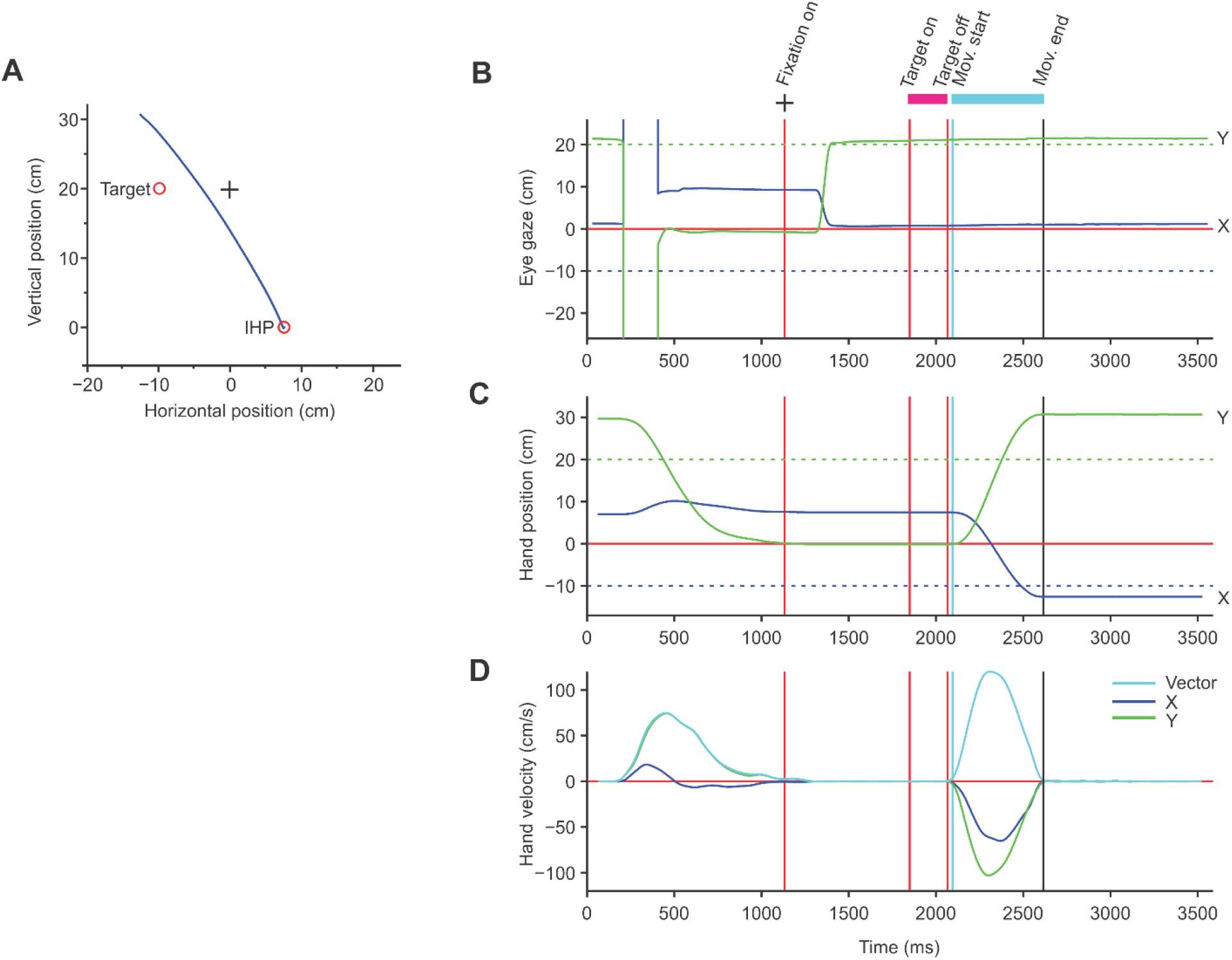
Typical participant trial. (A) Raw reaching trajectory from a typical trial. The required reach begins at the initial hand position, IHP, and ends at the target (red open circles). The blue line represents the participant’s hand trajectory. In this trial, the participant starts the reach with an IHP offset to the right by 7.5 cm. (B) Eye gaze traces. Horizontal (blue) and vertical (green) eye positions, from EyeLink recordings. Horizontal and vertical target position (dotted lines) are also plotted over time. Participants were instructed to keep the eyes fixated on the fixation cross (located at 0, 20 cm) throughout the trial. (C) Hand position traces. Horizontal (blue) and vertical (green) hand positions (solid lines) and horizontal and vertical target position (dotted lines) plotted over time. The trial begins when the subject aligns the fingertip indicator with the visual IHP and the simultaneous onset of the fixation cross. The fixation cross remains on until the end of the trial. The target then appears briefly. Participants were instructed to move in response to the flash of the target. Movement start and end times are represented by teal and black lines. (D) Hand velocity traces. Horizontal (blue), vertical (green), and vector (teal) hand velocities plotted over time.

### Data analyses

Offline analyses were performed using MATLAB (The Mathworks, Natick, MA). We smoothed the movement data using a low-pass, auto-regressive forward-backward filter (cutoff of 50Hz, eye data; 25Hz, hand). We removed trials in which participants broke fixation (Figure 5B).

A threshold of 1cm/s determined the onset/completion of a reach. The horizontal and vertical endpoint error were defined as the horizontal and vertical components of the displacement of the reach endpoint with respect to the target location.

Repeated measures ANOVAs were performed on changes in horizontal and vertical endpoint error, with the 4 factors, target location (−10, −5, 5, 10 cm), IHP (−7.5, 7.5 cm), polarity (anodal, cathodal), and site of stimulation (left mIPS, left PMd) at each epoch (change from baseline, during and after stimulation). ANOVA statistics were corrected for inhomogeneity of variance using the Greenhouse-Geisser correction when the Mauchly’s Test of Sphericity indicated that the assumption of sphericity had been violated. Three-way interactions were examined at each level of the fourth variable (site of stimulation), followed by two-way interactions at each level of the third variable (polarity). Interaction effects of IHP x target position, and/or main effects would indicate to us how the mIPS and PMd differ in their level of movement coding.

## Results

Only the cathodal polarity and post-stimulation epoch combination produced noticeable effects on horizontal endpoint error. Here, we focused on changes in horizontal endpoint error when reaching toward targets from the left and right IHPs after cathodal mIPS and PMd stimulation. Although the arrangement of the targets in the reaching task was designed to examine horizontal endpoint errors, vertical endpoint patterns also conformed to our interpretations (Figure S1, Appendix).

### Individual Participant Endpoint Error Analysis

In individual participants (Figures 6 and 7), the change in reach endpoint pattern following mIPS stimulation (gaze-centered expansion or contraction of reach endpoints) depended on the IHP (left or right), while the endpoint pattern was IHP-independent following PMd stimulation (contraction of reach endpoints for both IHP).

**Figure 6:**
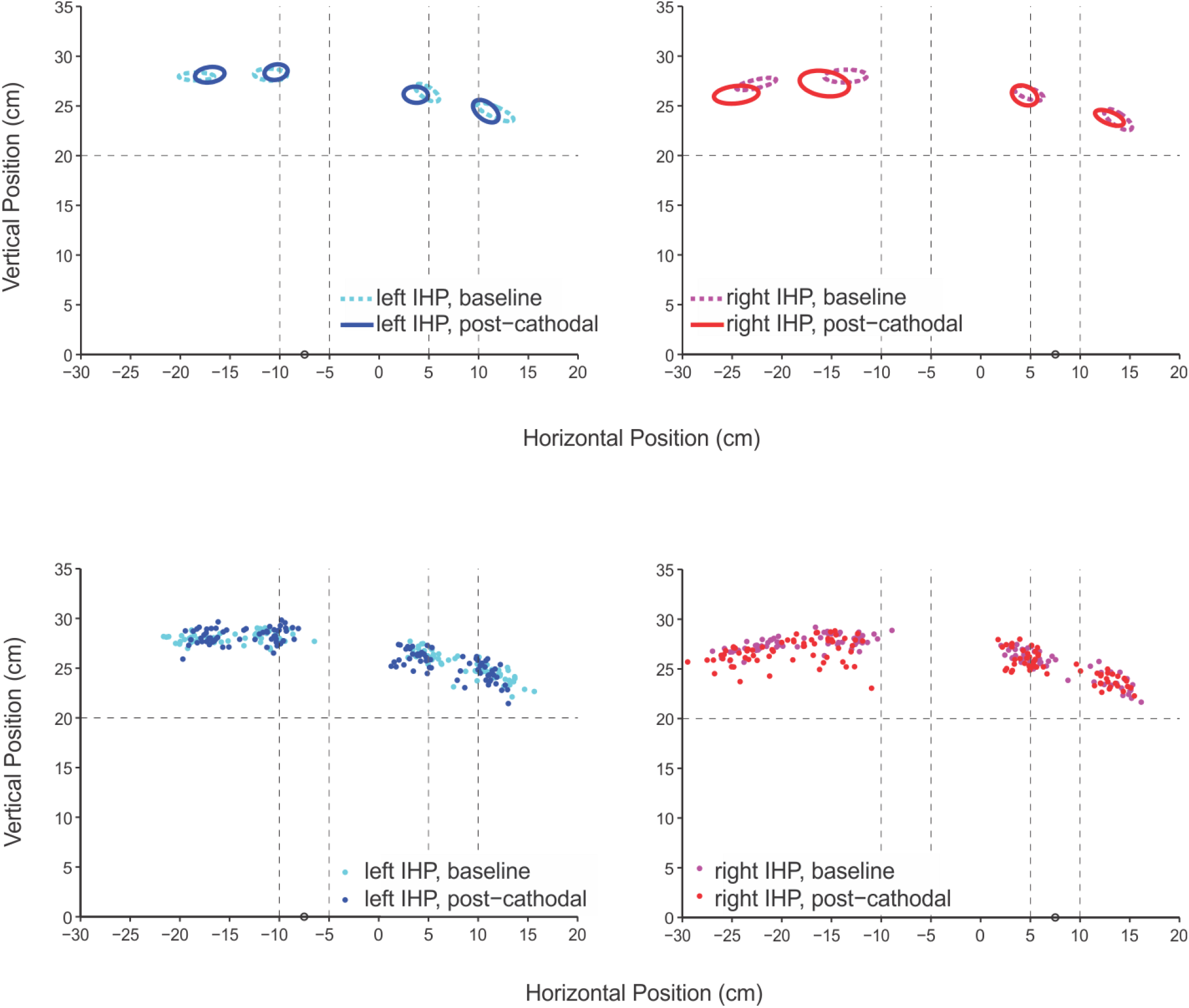
Reach endpoints pre- and post-cathodal mIPS stimulation. Standard ellipses of reach endpoints for each target location, with individual reach endpoints shown below, for all trials before (dotted teal and magenta) and after cathodal mIPS stimulation (solid blue and red), separated into reaches starting from the left (blues) and right IHP (reds), from a participant. The hand indicates the IHP and the intersections of the dashed guidelines represent target locations. It appears that there is a horizontal contraction effect of stimulation on the endpoints for reaches starting from the left IHP, but a horizontal expansion effect on the endpoints for reaches starting from the right IHP.

**Figure 7:**
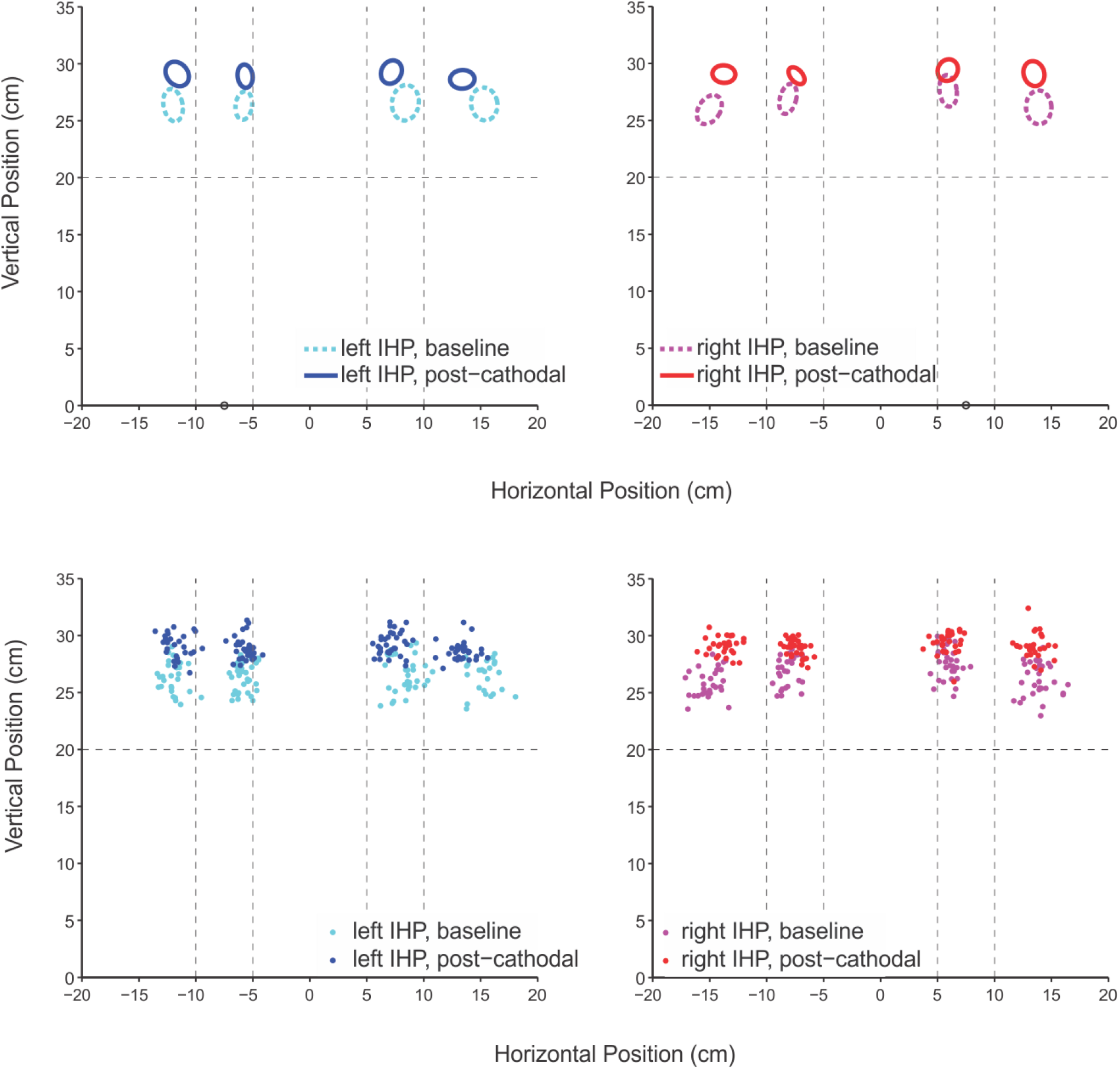
Reach endpoints pre- and post-cathodal PMd stimulation. Standard ellipses of reach endpoints for each target location, with individual reach endpoints shown below, for all trials before (dotted teal and magenta) and after cathodal PMd stimulation (solid blue and red), separated into reaches starting from the left (blues) and right IHP (reds), from a participant. The hand indicates the IHP and the intersections of the dashed guidelines represent target locations. It appears that there is a horizontal contraction effect of stimulation on the endpoints for reaches starting from either IHPs.

### Group-level Endpoint Error Analysis

Two separate two-way repeated measures ANOVAs at the group-level with the within-subject factors of IHP and target position on the change in horizontal error post vs pre-cathodal stimulation for each stimulation site were conducted. The full 4-way RM-ANOVA with the within-participant factors IHP, target position, polarity, and site revealed significant main target effect (F(3, 288) = 9.46, p = .0002), target*site interaction effect (F(3, 288) = 3.94, p = .0188), target*IHP*polarity interaction effect (F(3, 288) = 3.36, p = .0333), and IHP*polarity*site interaction effect (F(1, 288) = 31.2, p = .0003). We followed up the significant interactions consisting of the site term using separate 3-way RM-ANOVAs at each level of site, with the within-participant factors, polarity, target, and IHP. We found significant target*IHP*polarity interaction effect at the mIPS (F(3,144) = 4.26, p = .0138), and significant target main effect (F(3,144) = 13.8, p < .0001) and IHP*polarity interaction effect (F(1,144) = 4.07, p = .0288) at the PMd. We then followed up the significant interaction effects by conducting separate 2-way RM-ANOVAs at each level of polarity. The significant interaction and main effects are illustrated in Figure 8. An RM-ANOVA of the left mIPS post-cathodal change in horizontal endpoint error yielded a target x IHP interaction effect, *F*(3, 72) =3.31, *p* = .0350. A significant IHP by target position interaction effect demonstrates that the pattern across target positions differ according to IHP conditions (starting from the left or right) and would support separate IHP and target position coding at the input of mIPS. Contrastingly, at the left PMd, the RM-ANOVA yielded separate main effects of IHP, *F*(1, 72) = 7.7, *p* = .0216 and target position, *F*(3, 72) = 7, *p* = .0012, which support IHP-independent modulation of the movement vector and therefore movement vector coding at the input of PMd.

**Figure 8:**
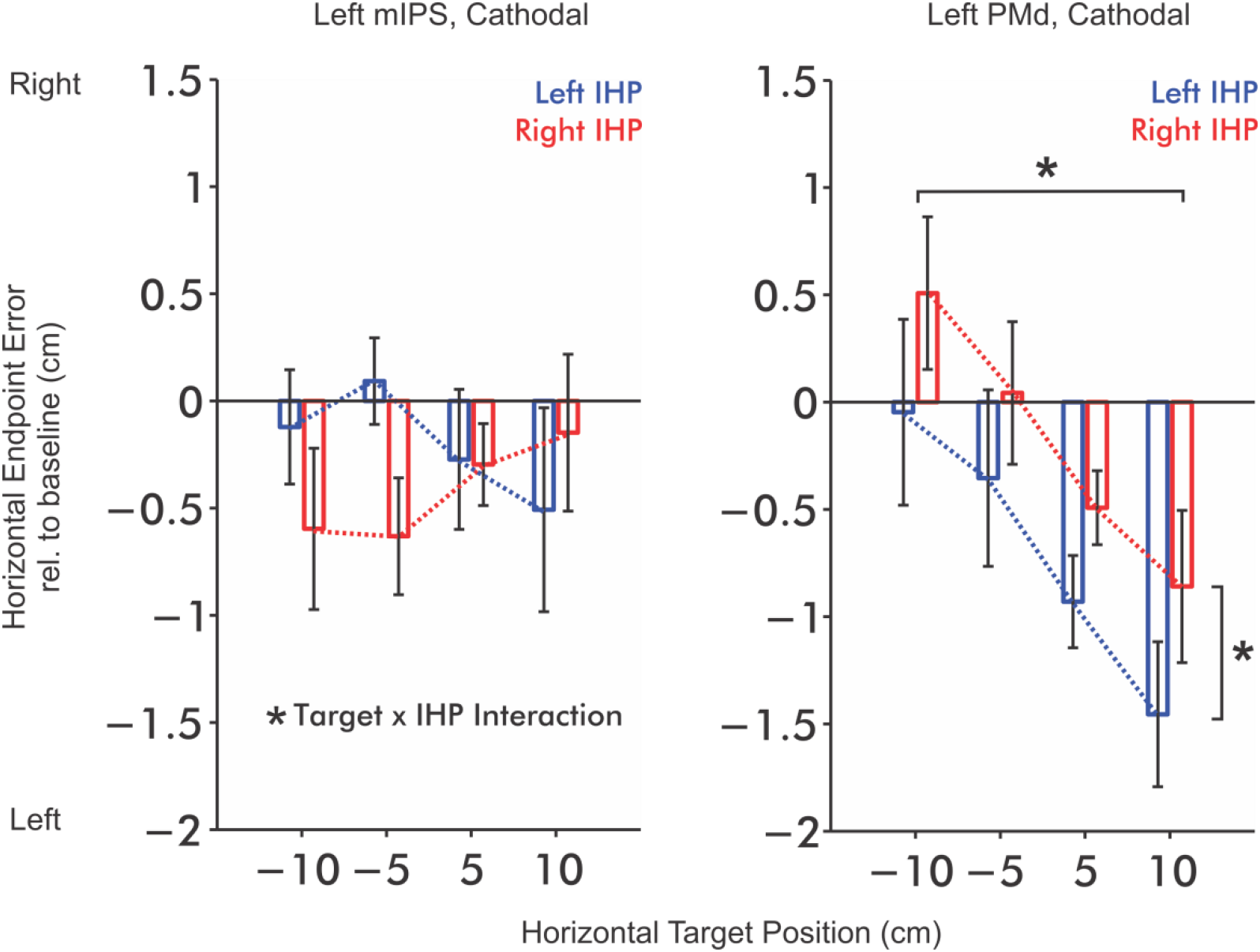
Group-level post-cathodal tDCS change in the horizontal endpoint error relative to baseline. Change in the horizontal component of reaching endpoint across target positions after cathodal stimulation of the left mIPS (left) and left PMd (right), separated according to left IHP (blue) and right IHP (red). Positive values represent endpoint falling to the right of the target and negative values represent endpoints falling to the left of the target. Error bars represent SEM. RM-ANOVAs of the post-cathodal change in horizontal endpoint error yielded a significant target x IHP interaction effect (* indicates p < .05) at the mIPS, suggesting independent IHP and target coding at the input of mIPS. Conversely, significant IHP and target main effects found at the PMd suggest that IHP and target position may be integrated into a movement vector code at the input of PMd.

**Figure 9:**
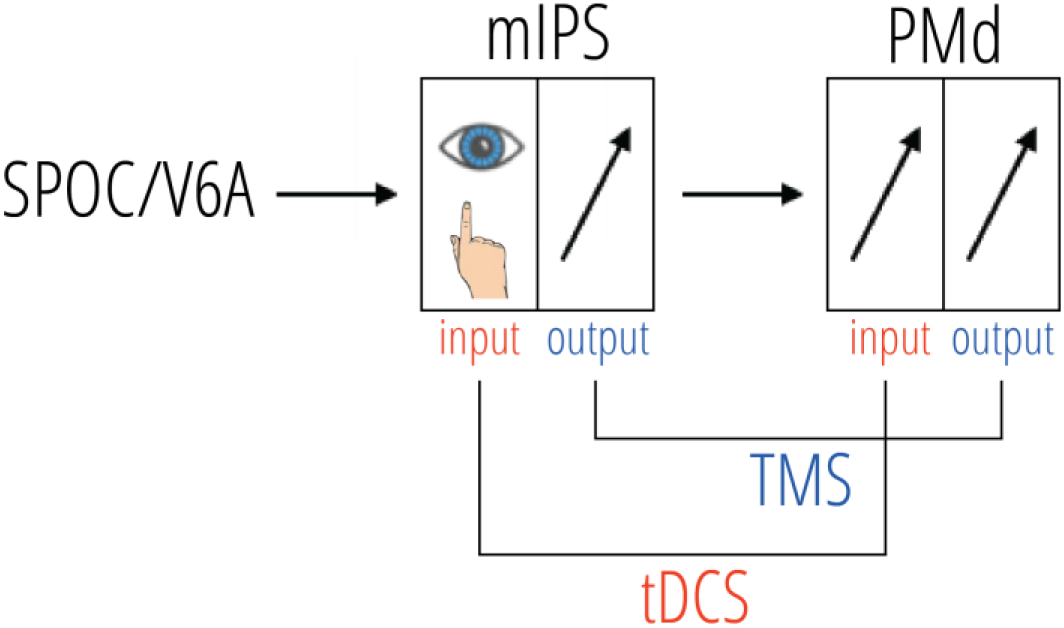
The model of the dorsomedial pathway we propose illustrates that the representation of the target position and IHP are separate at the input of the mIPS but are integrated into a vector representation at the input of the PMd.

### Alternative Explanations

We considered alternative explanations such as differences in information processing, speed-accuracy trade-offs, and feedback processes that could otherwise explain our results. Reaction and movement times did not significantly differ between conditions or between sessions (all *p*s > .05), therefore precluding the likelihood of our results being driven by differences in information processing and speed-accuracy trade-offs in movement execution. We did not find evidence of course adjustment during the movement from the hand position and velocity profiles (Figures S2 & S3, Appendix). Spatial trajectories were typically straight without terminal hooks and average velocity profiles were symmetrical and single peaked without additional valleys or inflections (Figure S4, Appendix), therefore precluding the likelihood of corrective sub-movements. This is not unusual given the lack of visual feedback. Thus, we do not have evidence to suggest that the changes in error patterns resulted from disturbed online control.

## Discussion

The current study is the first to apply HD-tDCS to directly investigate movement planning properties of the human mIPS and PMd. We found evidence that the mIPS receives independently coded target position and IHP information while the input of the PMd receives a vector-based movement plan. The present study demonstrates that the input of mIPS and PMd receive movement-relevant information at different levels of complexity towards the construction of a movement plan. Furthermore, we provide evidence that HD-tDCS can modulate performance in a movement planning task to support the understanding of the functional properties of mIPS and PMd.

Differences in horizontal error patterns were elicited from cathodal stimulation while that elicited from anodal stimulation were not significant. The mechanism that would explain the changes in behaviour resulting from cathodal stimulation is unlikely to be as straightforward as the general assumption of polarity-dependent modulation of excitability under the active electrode. Such an explanation would overlook the influence of the electrical field orientation across the convoluted cortical surface (Garnett, Malyutina, Datta, & den Ouden, 2015) which determines current direction with respect to the neuronal alignment and therefore the effect on neuronal excitability (Kabakov, Muller, Pascual-Leone, Jensen, & Rotenberg, 2012). Although the mechanism connecting excitability changes to behavioural changes has yet to be identified, the observable perturbations in movement errors permitted us to recognize how movement related components might be encoded at the input of the perturbed cortical areas.

The significant changes in movement errors produced by cathodal HD-tDCS in the stimulation offset period which were absent during stimulation lines up with the timeline of peak effects of the 4 x 1 HD-tDCS reported elsewhere, appearing not immediately after stimulation but half an hour after the end of stimulation (Villamar et al., 2013a; Villamar et al., 2013b; Kuo et al., 2013).

We observed behavioral changes in response to targeted HD-tDCS at the mIPS and PMd that would be consistent with independently coded target position and IHP by the former and dependent coding by the latter. The implications of this study for the use of HD-tDCS as a research technique is that there is a positive outlook for its use to clarify structure-function relationships in other brain areas and cognitive domains. The value of HD-tDCS lies in the focality that make it possible to target specific cortical areas. The validity of conclusions derived from this technique could be further strengthened by a complete mechanistic understanding of HD-tDCS.

In this study, we used a novel technique that distinguished the property of signals carried at the input of mIPS and PMd. This sets apart our experiment from TMS studies of the same areas where the effects are on the cortical output and processing at sites receiving projections from the stimulated site (Siebner, Hartwigsen, Kassuba, & Rothwell, 2009), whereas tDCS is thought to disrupt the input of a stimulated area, specifically, the presynaptic activity of stimulated neurons (Bączyk & Jankowska, 2014).

Together, the results of the two techniques assembles an overall picture of vector formation. Separably-encoded representations of the IHP and target position are provided at the input of mIPS and combined prior to the output of the mIPS. Together the present study and TMS literature closes in on mIPS as a location where vector formation occurs. For that reason, our results are complementary to rather than at odds with previous literature on the mIPS that proposed vector-based representation of the movement plan in the mIPS (Davare, Zénon, Desmurget, & Olivier, 2015; Vesia, Prime, Yan, Sergio, & Crawford, 2010). Our results about the PMd are also compatible with monkey neurophysiological and TMS findings that support a vector-based movement plan in the PMd (Kurata, 1993; Fu, Flament, Coltz, & Ebner, 1995; Davare, Zénon, Desmurget, & Olivier, 2015).

## Conclusion

Our results provide a plausible picture of where and how spatial information about the target position and IHP may be incorporated into a movement plan. The continued use and scientific acceptance of this exploratory tool for investigating the specific function of brain areas would benefit from identification of the mechanism in which HD-tDCS operates.

## Acknowledgements

SX & GB were supported by a NSERC Discovery Grant (Canada), a NSERC CREATE International Research and Training grant (Canada), and the Canadian Foundation for Innovation (CFI). JG received funding from NSERC Discovery Grant (Canada) and CFI award.

## Appendix

**Figure S1:**
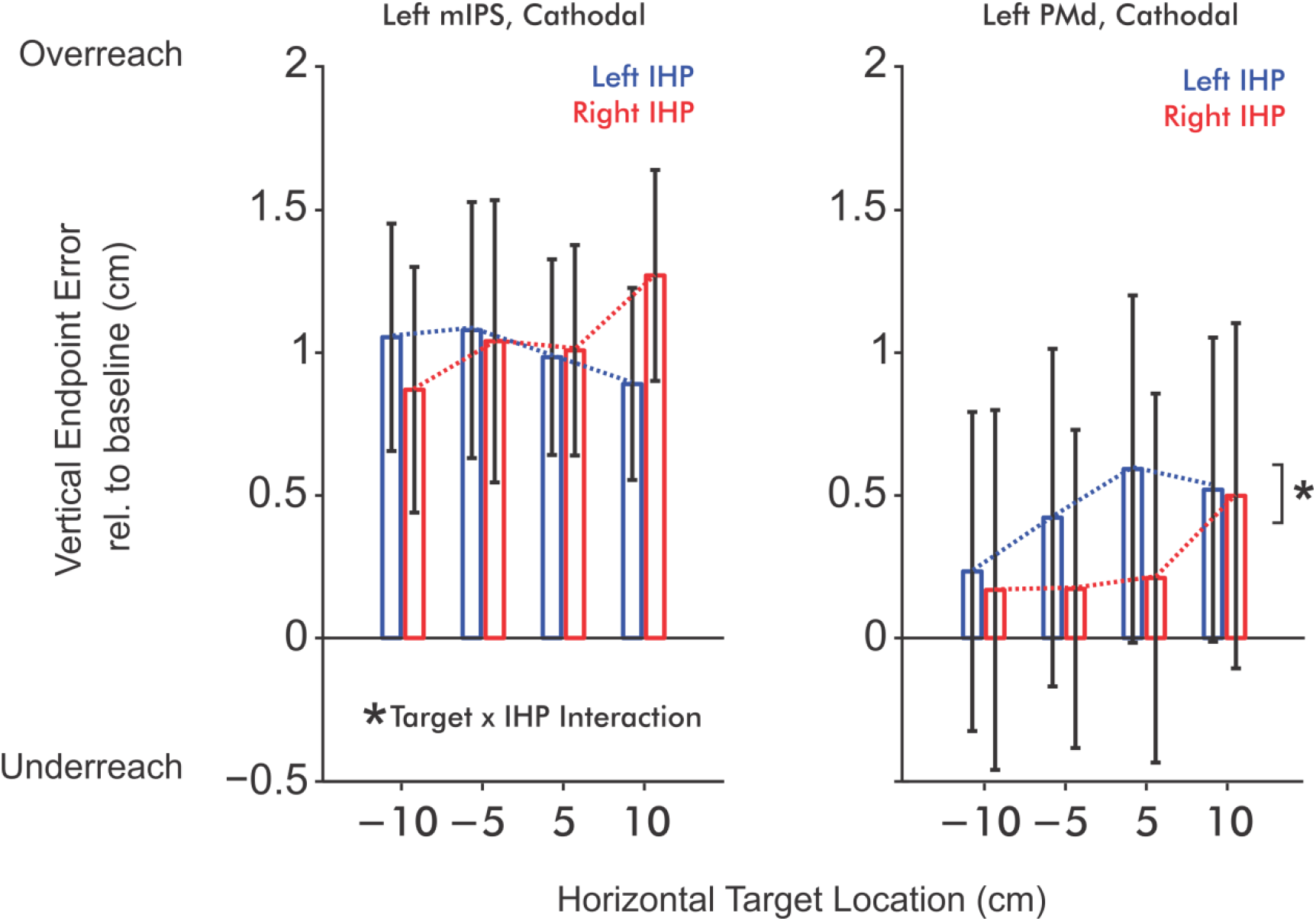
Group-level post-cathodal tDCS change in the vertical endpoint error relative to baseline. Change in the vertical component of reaching endpoint across target positions after cathodal stimulation of the left mIPS (left) and left PMd (right), separated according to left IHP (blue) and right IHP (red). Positive values indicate that the endpoint falls relative to the target, farther away from the participant while negative values, closer to participant. Error bars represent SEM. Similar to the results of horizontal endpoint error, RM-ANOVA yielded a significant target by IHP interaction (* indicates p < .05) at the mIPS. However, only a IHP main effect was found at the left PMd.

**Figure S2:**
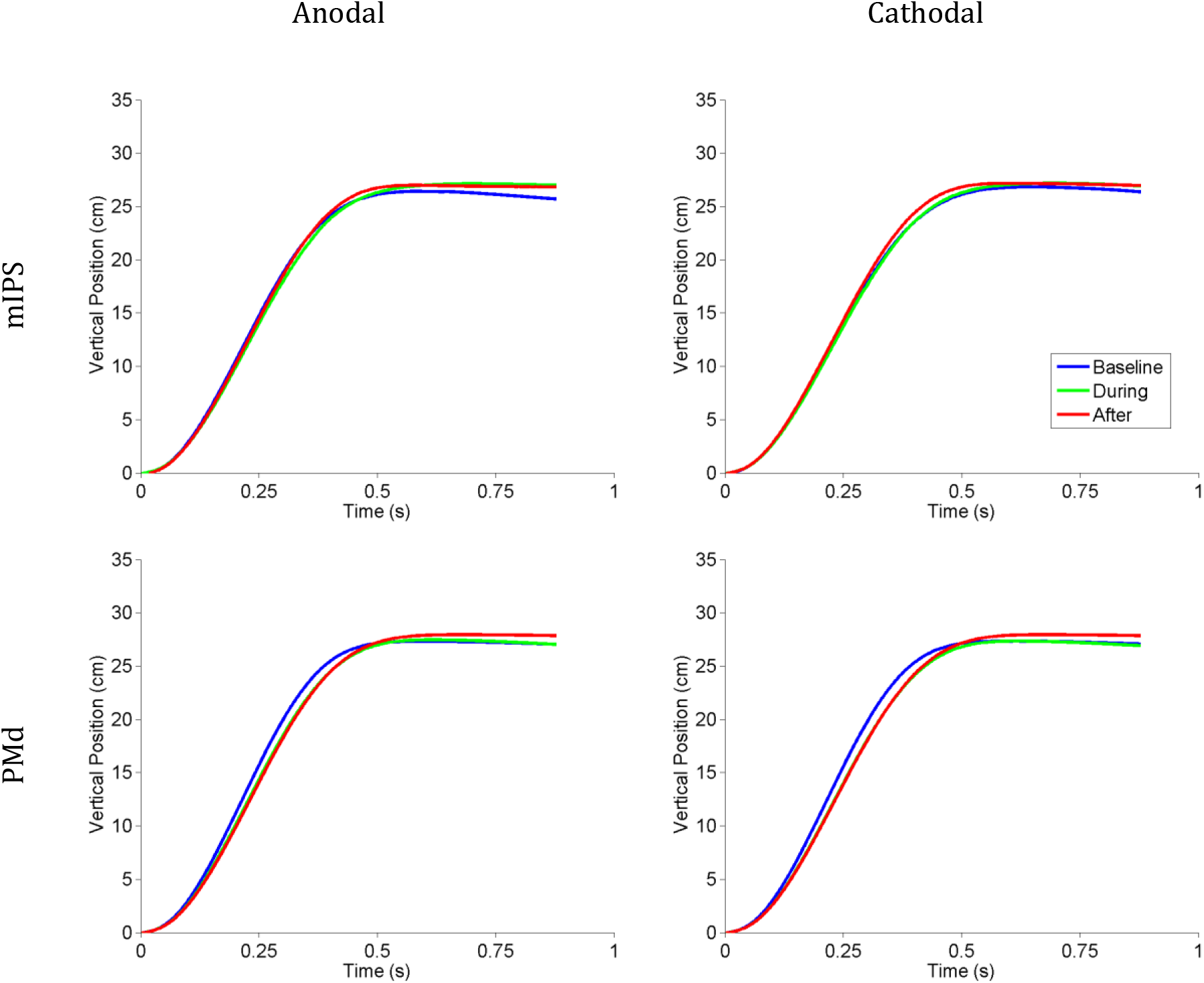
Average reaching position profiles. Average position along the near-far axis for a representative participant, for equal distance reaches (towards targets located in the same hemifield as the IHP), before, during and after anodal and cathodal mIPS and PMd tDCS. The position profiles along the near-far axis were nearly identical across conditions.

**Figure S3:**
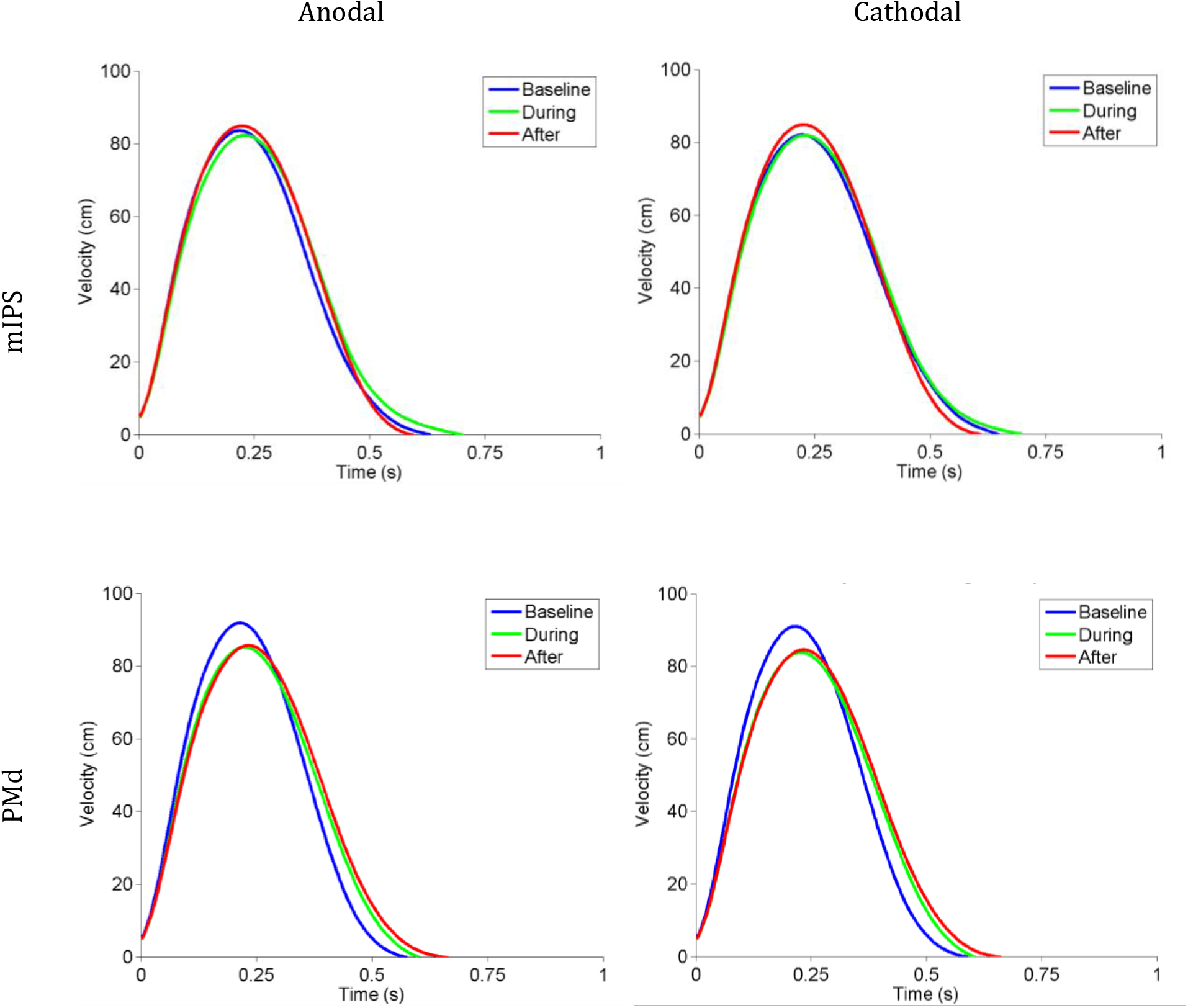
Average velocity profiles. Average hand velocities of a typical participant for equal distance reaches (towards targets located in the same hemifield as the IHP), before, during and after anodal and cathodal mIPS and PMd tDCS. The velocity profiles along the near-far axis were nearly identical across conditions.

**Figure S4:**
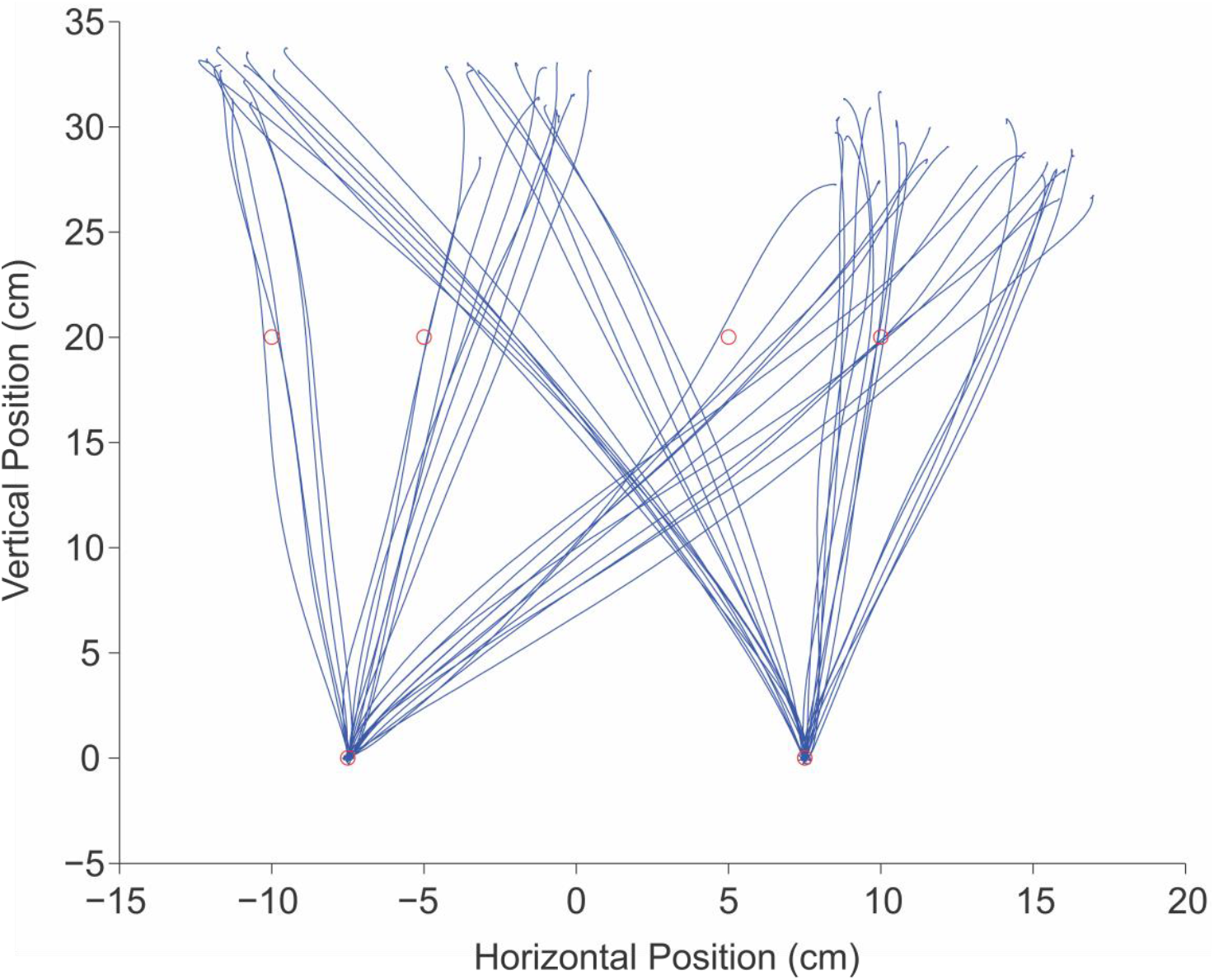
Reaching trajectories. Baseline trajectories made by an exemplary participant (P3) towards all targets starting from both IHPs for one block of trials, shown in blue lines. Red circles indicate starting hand positions (−7.5, 7.5 cm from midline) and target positions (−10, −5, 5, 10 cm from midline, 20 cm distant).

**Figure S5:**
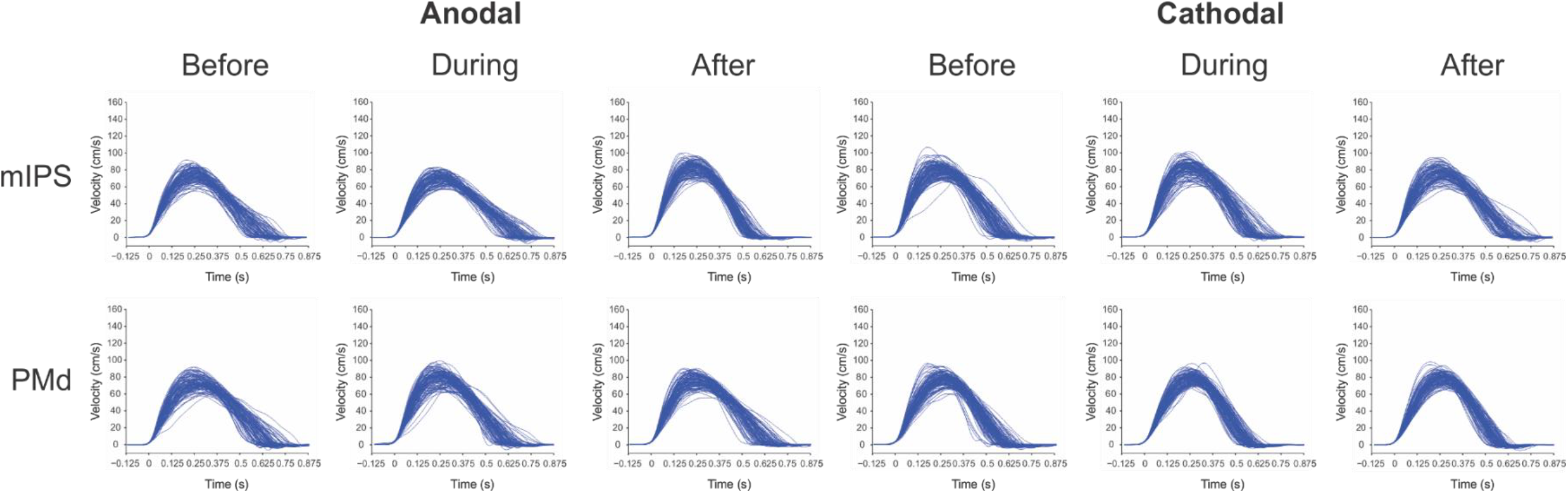
Velocity profiles of a representative participant. Velocity profiles before, during, and after anodal and cathodal tDCS of mIPS and PMd. Each line represents the velocity over time for a single trial. No additional valleys or inflections were observed in the individual trials to indicate the occurrence of sub-movements.

**Table S1.**
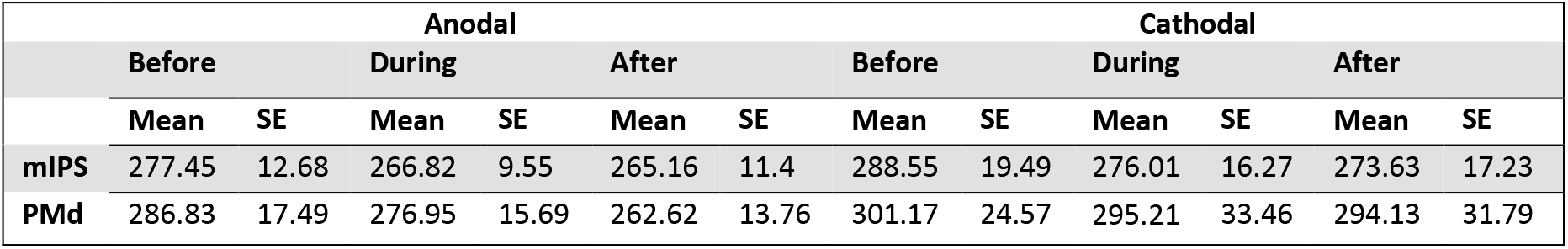
Mean reaction time (+/- SE). Mean reaction times at baseline, during and after anodal and cathodal mIPS and PMd stimulation epochs.

|  | Anodal |  |  |  |  |  | Cathodal |  |  |  |  |  |
| --- | --- | --- | --- | --- | --- | --- | --- | --- | --- | --- | --- | --- |
|  | Before |  | During |  | After |  | Before |  | During |  | After |  |
|  | Mean | SE | Mean | SE | Mean | SE | Mean | SE | Mean | SE | Mean | SE |
| **mIPS** | 277.45 | 12.68 | 266.82 | 9.55 | 265.16 | 11.4 | 288.55 | 19.49 | 276.01 | 16.27 | 273.63 | 17.23 |
| **PMd** | 286.83 | 17.49 | 276.95 | 15.69 | 262.62 | 13.76 | 301.17 | 24.57 | 295.21 | 33.46 | 294.13 | 31.79 |

**Table S2.** Mean movement time (+/- SE). Mean movement time at baseline, during and after anodal and cathodal mIPS and PMd stimulation epochs.

|  | Anodal |  |  |  |  |  | Cathodal |  |  |  |  |  |
| --- | --- | --- | --- | --- | --- | --- | --- | --- | --- | --- | --- | --- |
|  | Before |  | During |  | After |  | Before |  | During |  | After |  |
|  | Mean | SE | Mean | SE | Mean | SE | Mean | SE | Mean | SE | Mean | SE |
| **mIPS** | 613.9 | 39.34 | 582.4 | 28.35 | 560.4 | 22.05 | 587.5 | 25.28 | 583.0 | 25.94 | 587.1 | 27.16 |
| **PMd** | 583.2 | 32.85 | 588.45 | 27.95 | 582.5 | 26.48 | 598.0 | 30.46 | 587.9 | 22.92 | 575.3 | 21.13 |

**Table S3:**
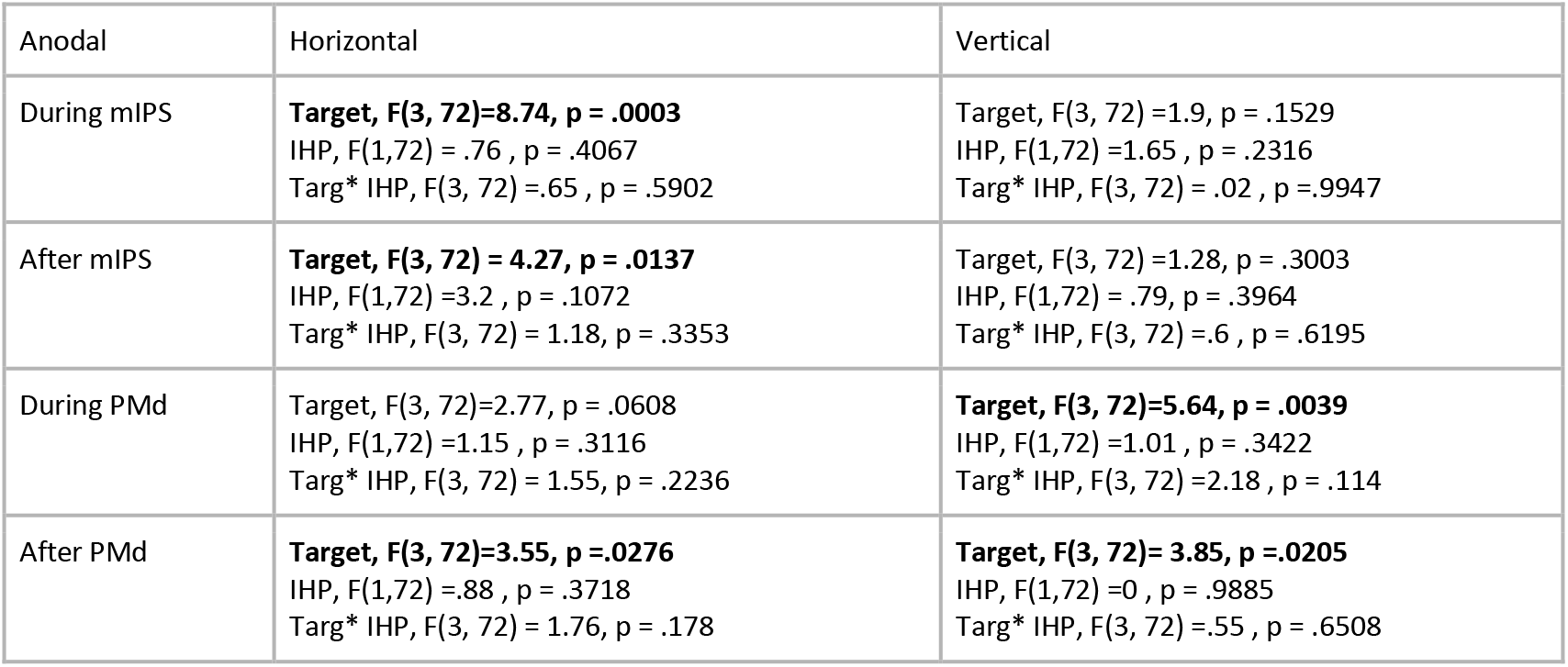
Results of 2-way RM-ANOVAs on the change in horizontal and vertical endpoint error, during and after anodal tDCS relative to baseline. Bolded represents significant effects at p < .05.

**Table S4:** Results of 2-way RM-ANOVAs on the change in horizontal and vertical endpoint error, during and after cathodal tDCS relative to baseline. Bolded represents significant effects at p < .05.

| Cathodal | Horizontal Error | Vertical Error |
| --- | --- | --- |
| During mIPS | Target, $F(3, 72) = .06, p = .982$<br>IHP, $F(1,72) = 1.78, p = .2152$<br>Targ* IHP, $F(3, 72) = 2.25, p = .1056$ | Target, $F(3, 72)=.37, p = .7737$<br>IHP, $F(1,72) = .35, p = .566$<br>Targ* IHP, $F(3, 72) = 1.51, p = .2351$ |
| After mIPS | Target, $F(3, 72)=.02, p = .9955$<br>IHP, $F(1,72) = 1.2, p = .3015$<br>**Targ\* IHP, $F(3, 72) = 3.31, p = .035$** | Target, $F(3, 72) = .27, p = .8457$<br>IHP, $F(1,72) = .12, p = .7399$<br>**Targ\* IHP, $F(3, 72) = 3.92, p = .0192$** |
| During PMd | **Target, $F(3, 72)=3.84, p = .0207$**<br>**IHP, $F(1,72) = 6.55, p = .0307$**<br>Targ* IHP, $F(3, 72) = .63, p = .6042$ | Target, $F(3, 72)= 2.93, p = .0516$<br>**IHP, $F(1,72) = 6.58, p = .0304$**<br>Targ* IHP, $F(3, 72) = 1.55, p = .2255$ |
| After PMd | **Target, $F(3, 72) = 7, p = .0012$**<br>**IHP, $F(1, 72) = 7.7, p = .0216$**<br>Targ * IHP, $F(3, 72) = .04, p = .8619$ | Target, $F(3, 72)= 1.86, p = .1605$<br>**IHP, $F(1,72) = 8.04, p = .0195$**<br>Targ* IHP, $F(3, 72) = .98, p = .4175$ |

**Table S5:** Curvature parameters during and after anodal and cathodal tDCS relative to baseline. CAR = the curvature-to-amplitude ratio, DIR = initial movement direction (at 200 ms), and θ, the angle at maximum curvature. None significant at alpha of .05.

| Anodal | CAR | DIR | $\theta$ |
| --- | --- | --- | --- |
| **mIPS, during** | Target, $F(3, 72) = .2, p = .894$<br>IHP, $F(1, 72) = .18, p = .6802$<br>Targ* IHP, $F(3, 72) = .17, p = .9155$ | Target, $F(3, 72) = .05, p = .985$<br>IHP, $F(1, 72) = .06, p = .8146$<br>Targ* IHP, $F(3, 72) = .02, p = .9957$ | Target, $F(3, 72) = .05, p = .9833$<br>IHP, $F(1, 72) = 0, p = .996$<br>Targ* IHP, $F(3, 72) = .53, p = .6675$ |
| **mIPS, after** | Target, $F(3, 72) = 1.41, p = .2626$<br>IHP, $F(1, 72) = .2, p = .669$<br>Targ* IHP, $F(3, 72) = .26, p = .8506$ | Target, $F(3, 72) = .29, p = .8341$<br>IHP, $F(1, 72) = .27, p = .6127$<br>Targ* IHP, $F(3, 72) = .36, p = .7805$ | Target, $F(3, 72) = .53, p = .6653$<br>IHP, $F(1, 72) = 0, p = .9986$<br>Targ* IHP, $F(3, 72) = .6, p = .6235$ |
| **PMd, during** | Target, $F(3, 72) = 1.15, p = .3451$<br>IHP, $F(1, 72) = 1.76, p = .2174$<br>Targ* IHP, $F(3, 72) = .65, p = .5899$ | Target, $F(3, 72) = .73, p = .5404$<br>IHP, $F(1, 72) = 1.82, p = .2106$<br>Targ* IHP, $F(3, 72) = 2.32, p = .0977$ | Target, $F(3, 72) = 2.12, p = .1211$<br>IHP, $F(1, 72) = .43, p = .5283$<br>Targ* IHP, $F(3, 72) = .78, p = .513$ |
| **PMd, after** | Target, $F(3, 72) = 1.37, p = .274$<br>IHP, $F(1, 72) = 1.24, p = .2951$<br>Targ* IHP, $F(3, 72) = .86, p = .4732$ | Target, $F(3, 72) = .7, p = .5596$<br>IHP, $F(1, 72) = .31, p = .5893$<br>Targ* IHP, $F(3, 72) = .65, p = .5926$ | Target, $F(3, 72) = 1.06, p = .3831$<br>IHP, $F(1, 72) = .49, p = .5031$<br>Targ* IHP, $F(3, 72) = .8, p = .5035$ |
| **Cathodal** | **CAR** | **DIR** | **$\theta$** |
| **mIPS, during** | Target, $F(3, 72) = 1.24, p = .3154$<br>IHP, $F(1, 72) = 0, p = .99446$<br>Targ* IHP, $F(3, 72) = 2.46, p = .0844$ | Target, $F(3, 72) = 1.57, p = .2195$<br>IHP, $F(1, 72) = .09, p = .7724$<br>Targ* IHP, $F(3, 72) = 1.37, p = .2737$ | Target, $F(3, 72) = 1.83, p = .1648$<br>IHP, $F(1, 72) = .11, p = .7442$<br>Targ* IHP, $F(3, 72) = 2.73, p = .0634$ |
| **mIPS, after** | Target, $F(3, 72) = .6, p = .6189$<br>IHP, $F(1, 72) = .15, p = .7051$<br>Targ* IHP, $F(3, 72) = 2.32, p = .0982$ | Target, $F(3, 72) = .69, p = .568$<br>IHP, $F(1, 72) = .32, p = .5878$<br>Targ* IHP, $F(3, 72) = 2.64, p = .07$ | Target, $F(3, 72) = .73, p = .5438$<br>IHP, $F(1, 72) = .11, p = .7442$<br>Targ* IHP, $F(3, 72) = 2.83, p = .0569$ |
| **PMd, during** | Target, $F(3, 72) = 1.71, p = .1886$<br>IHP, $F(1, 72) = .06, p = .8105$<br>Targ* IHP, $F(3, 72) = .61, p = .6149$ | Target, $F(3, 72) = 1.5, p = .2377$<br>IHP, $F(1, 72) = .93, p = .3603$<br>Targ* IHP, $F(3, 72) = .04, p = .9886$ | Target, $F(3, 72) = 2.49, p = .0818$<br>IHP, $F(1, 72) = 2.28, p = .165$<br>Targ* IHP, $F(3, 72) = .26, p = .8551$ |
| **PMd, after** | Target, $F(3, 72) = 1.61, p = .2109$<br>IHP, $F(1, 72) = .02, p = .8829$<br>Targ* IHP, $F(3, 72) = .48, p = .6984$ | Target, $F(3, 72) = 2.39, p = .0903$<br>IHP, $F(1, 72) = .32, p = .5849$<br>Targ* IHP, $F(3, 72) = .82, p = .4942$ | Target, $F(3, 72) = 2.72, p = .064$<br>IHP, $F(1, 72) = .2, p = .6632$<br>Targ* IHP, $F(3, 72) = .46, p = .7105$ |

**Table S6:** Comparisons between curvature parameters before, during, and after anodal and cathodal tDCS. CAR = the curvature-to-amplitude ratio, DIR = initial movement direction (at 200 ms), and θ, the angle at maximum curvature. None significant at alpha of .05.

| CAR | Comparison |  | Difference between estimated group means | Lower bound of 95% confidence interval for true mean difference | Upper bound of 95% confidence interval for true mean difference | p-value |
| --- | --- | --- | --- | --- | --- | --- |
|  | Pre-anodal | During | .0004 | -.0077 | .0085 | .9992 |
|  | Pre-anodal | Post-anodal | -.0007 | -.0087 | .0074 | .9964 |
|  | During | Post-anodal | -.0010 | -.0091 | .0070 | .9856 |
|  | Pre-cathodal | During | -.0052 | -.0157 | .0054 | .5647 |
|  | Pre-cathodal | Post-cathodal | -.0072 | -.0178 | .0033 | .2744 |
|  | During | Post-cathodal | -.0021 | -.0127 | .0085 | .9522 |

| DIR | Comparison |  | Difference between estimated group means | Lower bound of 95% confidence interval for true mean difference | Upper bound of 95% confidence interval for true mean difference | p-value |
| --- | --- | --- | --- | --- | --- | --- |
|  | Pre-anodal | During | -.0975 | -.3617 | .1667 | .7563 |
|  | Pre-anodal | Post-anodal | -.1878 | -.4520 | .0763 | .2426 |
|  | During | Post-anodal | -.0903 | -.3545 | .1738 | .7960 |
|  | Pre-cathodal | During | -.0975 | -.3617 | .1667 | .7563 |
|  | Pre-cathodal | Post-cathodal | -.1878 | -.4520 | .0763 | .2426 |
|  | During | Post-cathodal | -.0903 | -.3545 | .1738 | .7960 |

| $\theta$ | Comparison | | Difference between estimated group means | Lower bound of 95% confidence interval for true mean difference | Upper bound of 95% confidence interval for true mean difference | p-value |
| --- | --- | --- | --- | --- | --- | --- |
|  | Pre-anodal | During | -.7134 | -2.1305 | .7037 | .5381 |
|  | Pre-anodal | Post-anodal | -.9394 | -2.3564 | .4777 | .3000 |
|  | During | Post-anodal | -.2260 | -1.6430 | 1.1911 | .9734 |
|  | Pre-cathodal | During | -.7134 | -2.1305 | .7037 | .5381 |
|  | Pre-cathodal | Post-cathodal | -.9394 | -2.3564 | .4777 | .3000 |
|  | During | Post-cathodal | -.2260 | -1.6430 | 1.1911 | .9734 |

## References

Amiez, C., Kostopoulos, P., Champod, A.-S., & Petrides, M. (2006). Local morphology predicts functional organization of the dorsal premotor region in the human brain. The Journal of Neuroscience: The Official Journal of the Society for Neuroscience, 26(10), 2724–2731. https://doi.org/10.1523/JNEUROSCI.4739-05.2006

Bączyk, M., & Jankowska, E. (2014). Presynaptic actions of transcranial and local direct current stimulation in the red nucleus: Presynaptic modulation. The Journal of Physiology, 592(19), 4313–4328.

Batista, A. P., Buneo, C. A., Snyder, L. H., & Andersen, R. A. (1999). Reach plans in eye-centered coordinates. Science, 285(5425), 257–260.

Bindman, L. J., Lippold, O. C., & Redfearn, J. W. (1962). Long-lasting changes in the level of the electrical activity of the cerebral cortex produced by polarizing currents. Nature, 196, 584–585.

Blangero, A., Menz, M. M., McNamara, A., & Binkofski, F. (2009). Parietal modules for reaching. Neuropsychologia, 47(6), 1500–1507.

Bock, O., & Arnold, K. (1992). Motor control prior to movement onset: Preparatory mechanisms for pointing at visual targets. Experimental Brain Research, 90(1), 209–216.

Cisek, P. (2002). Neural Activity in Primary Motor and Dorsal Premotor Cortex In Reaching Tasks With the Contralateral Versus Ipsilateral Arm. Journal of Neurophysiology, 89(2), 922–942.

Datta, A., Bansal, V., Diaz, J., Patel, J., Reato, D., & Bikson, M. (2009). Gyri-precise head model of transcranial direct current stimulation: Improved spatial focality using a ring electrode versus conventional rectangular pad. Brain Stimulation, 2(4), 201–207.

Datta, A., Truong, D., Minhas, P., Parra, L. C., & Bikson, M. (2012). Inter-individual variation during transcranial direct current stimulation and normalization of dose using MRI derived computational models. Frontiers in Psychiatry, 3, 91.

Davare, M., Zénon, A., Desmurget, M., & Olivier, E. (2015). Dissociable contribution of the parietal and frontal cortex to coding movement direction and amplitude. Frontiers in Human Neuroscience, 9, 241.

Davare, M., Zenon, A., Pourtois, G., Desmurget, M., & Olivier, E. (2012). Role of the medial part of the intraparietal sulcus in implementing movement direction. Cerebral Cortex, 22(6), 1382–1394.

Fu, Q. G., Flament, D., Coltz, J. D., & Ebner, T. J. (1995). Temporal encoding of movement kinematics in the discharge of primate primary motor and premotor neurons. Journal of Neurophysiology, 73(2), 836–854.

Gallivan, J. P., McLean, D. A., Smith, F. W., & Culham, J. C. (2011). Decoding effector dependent and effector-independent movement intentions from human parieto-frontal brain activity. Journal of Neuroscience, 31(47), 17149–17168.

Grefkes, C., Ritzl, A., Zilles, K., & Fink, G. R. (2004). Human medial intraparietal cortex subserves visuomotor coordinate transformation. NeuroImage, 23(4), 1494–1506.

Jeannerod M, Arbib MA, Rizzolatti G, Sakata H. (1995). Grasping objects: the cortical mechanisms of visuomotor transformation. Trends Neuroscience, 18, 314–320.

Krakauer, J. W., Pine, Z. M., Ghilardi, M. F., & Ghez, C. (2000). Learning of visuomotor transformations for vectorial planning of reaching trajectories. The Journal of Neuroscience, 20(23), 8916–8924.

Kuo, H.-I., Bikson, M., Datta, A., Minhas, P., Paulus, W., Kuo, M.-F., & Nitsche, M. A. (2013). Comparing cortical plasticity induced by conventional and high-definition 4 Å~ 1 ring tDCS: A neurophysiological study. Brain Stimulation, 6(4), 644–648.

Kurata, K. (1993). Premotor cortex of monkeys: set-and movement-related activity reflecting amplitude and direction of wrist movements. Journal of Neurophysiology, 69(1), 187–200.

Mayka, M. A., Corcos, D. M., Leurgans, S. E., & Vaillancourt, D. E. (2006). Three-dimensional locations and boundaries of motor and premotor cortices as defined by functional brain imaging: a meta-analysis. NeuroImage, 31(4), 1453–1474.

Prado, J., Clavagnier, S., Otzenberger, H., Scheiber, C., Kennedy, H., & Perenin, M.-T. (2005). Two cortical systems for reaching in central and peripheral vision. Neuron, 48(5), 849–858.

Purpura, D. P., & Mcmurtry, J. G. (1965). Intracellular activities and evoked potential changes during polarization of motor cortex. Journal of Neurophysiology, 28, 166–185.

Siebner, H. R., Hartwigsen, G., Kassuba, T., & Rothwell, J. C. (2009). How does transcranial magnetic stimulation modify neuronal activity in the brain? Cortex, 45(9), 1035–1042.

Stark, A., & Zohary, E. (2008). Parietal mapping of visuomotor transformations during human tool grasping. Cerebral Cortex, 18(10), 2358–2368.

Vesia, M., Prime, S. L., Yan, X., Sergio, L. E., & Crawford, J. D. (2010). Specificity of human parietal saccade and reach regions during transcranial magnetic stimulation. Journal of Neuroscience, 30(39), 13053–13065.

Villamar, M. F., Volz, M. S., Bikson, M., Datta, A., Dasilva, A. F., & Fregni, F. (2013). Technique and considerations in the use of 4×1 ring high-definition transcranial direct current stimulation (HD-tDCS). Journal of Visualized Experiments: JoVE, (77), e50309.

Villamar, M. F., Wivatvongvana, P., Patumanond, J., Bikson, M., Truong, D. Q., Datta, A., & Fregni, F. (2013). Focal modulation of the primary motor cortex in fibromyalgia using 4Å~1-ring high-definition transcranial direct current stimulation (HD-tDCS): immediate and delayed analgesic effects of cathodal and anodal stimulation. The Journal of Pain, 14(4), 371–383.

Vindras, P., & Viviani, P. (1998). Frames of reference and control parameters in visuomanual pointing. Journal of Experimental Psychology, 24(2), 569–591.

Vingerhoets, G. (2014). Contribution of the posterior parietal cortex in reaching, grasping, and using objects and tools. Frontiers in Psychology, 5.

